# The Relationships of Resting-state Brain Entropy (BEN), Ovarian Hormones and Behavioral Inhibition and Activation Systems (BIS/BAS)

**DOI:** 10.1101/2024.06.04.595915

**Authors:** Dong-Hui Song, Ze Wang

**Affiliations:** State Key Laboratory of Cognitive Neuroscience and Learning, Beijing Normal University, Beijing 100091, China; IDG/McGovern Institute for Brain Research, Beijing Normal University, Beijing 100091, China; Department of Diagnostic Radiology and Nuclear Medicine, University of Maryland School of Medicine, Baltimore, MD, 21201, USA

**Keywords:** Resting-state fMRI, Brain entropy (BEN), Hormones, Behavioral inhibition and activation systems, Mediation analysis

## Abstract

Entropy measures the irregularity or complexity of a system. Recent research on brain entropy (BEN) based on resting-state fMRI has provided complementary information to other metrics such as low-frequency fluctuations and cerebral blood flow. It has been established that neural plasticity, both pharmacological and nonpharmacological, as well as brain stimulation can influence BEN. However, it remains unknown whether BEN can reflect the effects of hormones. Furthermore, recent studies have indicated that ovarian hormones influence both the behavioral inhibition and activation systems. In our study, we utilized open-access available data from OpenNeuro to investigate the effects of ovarian hormones on BEN and their impact on BIS/BAS.

Our results indicated a negative correlation between progesterone (PROG) and BEN in the frontal-parietal network and limbic system, while BEN showed a significant positive correlation with BAS-drive in the DLPFC. Additionally, a significant negative correlation was observed between PROG and BAS-drive. Further analysis revealed that DLPFC BEN mediates the negative correlation between PROG and BAS-drive. This suggests that PROG reduces BAS-drive by increasing the executive and inhibitory functions of DLPFC. We also analyzed the FC between DLPFC and the whole brain. DLPFC-IPL FC showed a significant positive correlation with BAS-drive, while DLPFC-LOFC FC exhibited a significant negative correlation with BAS-fun-seeking. Moreover, DLPFC-AG FC demonstrated a significant positive correlation with BAS-rewards. These results are consistent with the relationship between executive functions of the frontal-parietal network and impulsivity representation of BAS.

Our study is the first to demonstrate that BEN can also reflect the impact of hormones on brain function. Additionally, we identified that the negative correlation between PROG and BAS-drive is mediated by left DLPFC BEN, providing new insights into our understanding of the effects of PROG on the brain and behavior.

## 1 Introduction

Entropy originally comes from thermodynamics, measuring the irregularity and disorders of a dynamic system [1], and entropy increases over time, ultimately reaching a state of maximum chaos and disorder in an isolated system, according to the second law of thermodynamics [2]. Subsequently, entropy quantifies uncertainty, surprise, and the amount of information in information theory [3] and has also been adapted for physiological signals [4]. As one of the most complex systems in the world, the human brain consumes a lot of energy [5] to reduce its entropy and uncertainty to maintain proper functioning through constant interaction with the external environment [6–9]. Therefore, entropy can serve as a metric to depict the intricate dynamic state of the brain, observing normal and disrupted brain function. In recent years, there has been increasing recognition of the value of brain entropy (BEN) derived from resting-state functional magnetic resonance imaging (rs-fMRI) [10], which offers additional complementary insights into spontaneous brain activity [11], alongside widely used measures such as the fractional amplitude of low-frequency fluctuations (fALFF) [12] and cerebral blood flow (CBF) [13–15]. As a result, interest in BEN is rapidly growing [10, 16, 17]. The relationship between BEN and cognitive function has been identified [10, 16, 18, 19], and mental disorders related to BEN patterns have also been found in a variety of brain diseases [20–26]. More importantly, BEN can reflect the effects of pharmacological [20, 27] and nonpharmacological treatments for depression [28]. While BEN can also be altered by non-invasive brain stimulation [29–31] in normal young adults, indicating that BEN is sensitive to neuroplasticity. However, it remains uncertain whether BEN can effectively reflect the influence of hormones on brain function, given that a considerable body of research has demonstrated the effects of hormones on brain structure and function [32–35]. Investigating the relationship between BEN and hormones can provide new insights into how these chemical molecules affect brain function. In this study, we utilized ovarian hormones to conduct a preliminary investigation into the association between BEN and hormones, aiming to address this knowledge gap.

Ovarian hormones are reported to brain structures and functions [32, 33, 36] including executive function [37–40], emotion processing [37, 41, 42], and reward [43, 44]. Progesterone (PROG) and Estradiol (E2) are two main ovarian hormones that regularly fluctuate during women’s menstrual cycles [45]. Generally, the menstrual cycle can be divided into two phases: the follicular phase (FP), between the onset of menses and ovulation with rising levels of E2 and deficient levels of PROG; the luteal phase (LP), starting after ovulation until the onset of the next menses characterized by high levels of PROG and the second rising levels of E2 in the mid-LP [45]. A recent study also showed that ovarian hormones influence the behavioral activation and inhibition system (BIS/BAS) [46]. BAS is associated with sensitivity to reward and approach motivation, while BIS is related to sensitivity to punishment and avoidance motivation [47, 48]. BIS/BAS is widely thought to be associated with affective disorders [49–52] and addiction [53–55]. As previously mentioned, ovarian hormones also exert an influence on emotion and reward systems. Our recent research on BEN indirectly suggests a potential association between ovarian hormones and BIS/BAS. We observed elevated BEN in the DLPFC and limbic system among patients with depression, indicating emotion dysregulation [28] and consistently higher BEN in the DLPFC among substance use disorders including nicotine smoking, marijuana use, and alcohol use [22]. Moreover, in adolescents, higher prefrontal-parietal BEN was associated with impulsivity and drug use risk [56]. Therefore, the relationship between BEN and BIS/BAS may also be affected by ovarian hormones.

In summary, the study first explores whether BEN is affected by ovarian hormones and then examines whether it subsequently affects the relationship between BEN and BIS/BAS.

## 2 Results

### 2.1 Quality Control and Demographic Information

Five participants were removed from the dataset, two of whom (sub-122, sub-230) had rs-fMRI with large head motion (mean FD > 0.2 mm), and three of whom (sub-117, sub-121, sub-238) had rs-fMRI with artifacts. Finally, forty-four participants (age=22.61±2.14 years) were included in the study, of which LP included 21 participants (age=22.76±1.84 years), and FP included 23 participants (age=22.48±2.41 years). There are no significant differences in age (*t*=0.44, *P*=0.665) and mean FD between FL and FP (*t*=1.72, *P*=0.093).

### 2.2 Hormones assays and BIS/BAS scoring

There are no significant differences in PROG, E2, and BIS/BAS for different menstrual phases (see Table 1).

**Table 1.** Hormone assays and BIS/BAS scoring for each menstrual phase.

| Variable | All (M±SD) | LP(M±SD) | FP(M±SD) | *t/p* |
| --- | --- | --- | --- | --- |
| PROG | 101.63±118.34 | 133.79±129.37 | 72.27±101.33 | 1.764/0.085 |
| E2 | 5.73±3.29 | 5.18±2.42 | 6.22±3.91 | -1.051/0.299 |
| BAS-drive | 13.86±1.50 | 13.81±1.21 | 13.91±1.76 | -0.226/0.823 |
| BAS-rewards | 12.57±1.90 | 12.52±2.06 | 12.61±1.78 | -0.147/0.884 |
| BAS-fun seeking | 14.91±2.25 | 14.95±2.33 | 14.87±2.22 | 0.121/0.904 |
| BIS | 16.36±2.33 | 16.43±1.99 | 16.3±2.65 | 0.174/0.862 |

Note: M±SD: Mean ± Standard Deviation, PROG: Progesterone, E2: Estradiol, ALL: All participants, LP: follicular phase group, FP: luteal phase group. *t/p*: *t* value/*p* value from two sample *t*-tests for the LP group and FP group.

### 2.3 The BEN differences between LP and FP

No significant BEN differences were found between LP and FP.

### 2.4 Correlation of BEN with Ovarian Hormones

The negative correlation between BEN and PROG was observed in the right amygdala, right hippocampus (HPC), parahippocampal cortex (PHPC), and dorsolateral prefrontal cortex (DLPFC) (Fig 1a). When E2 was added as a covariate, a significant negative correlation also emerged in the right inferior parietal lobule (IPL), right primary sensory cortex (PSC), right angular gyrus (AG), and right primary motor cortex (PMC) (Fig 1b). No significant correlation was observed between BEN and E2, even after the inclusion of PROG as a covariate. Table 2 provides information on clusters exceeding the threshold (voxel-wise p < 0.001, cluster size ≥ 810 mm^3^), including peak MNI coordinates, cluster size, label on the AAL atlas, and brain regions for correlation analysis of BEN with ovarian hormones.

**Fig 1.**
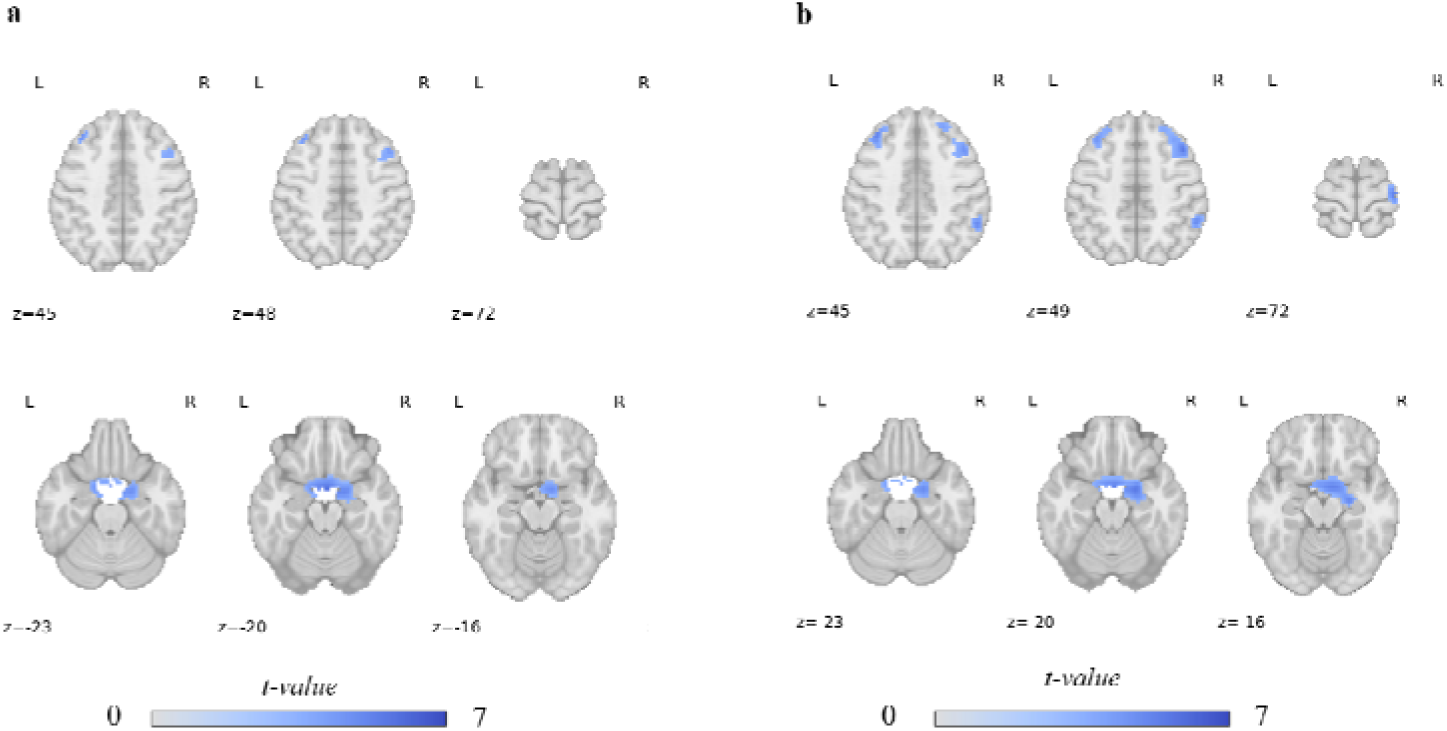
The correlations of BEN-PROG, (a) without E2 as a covariance, (b) add E2 as a covariance. The color bar indicates the t value, and the blue indicates a negative correlation between BEN and PROG. The number beneath each slice indicates its location along the z-axis in the MNI space, and L means left hemisphere, and R means right hemisphere.

**Table 2.**
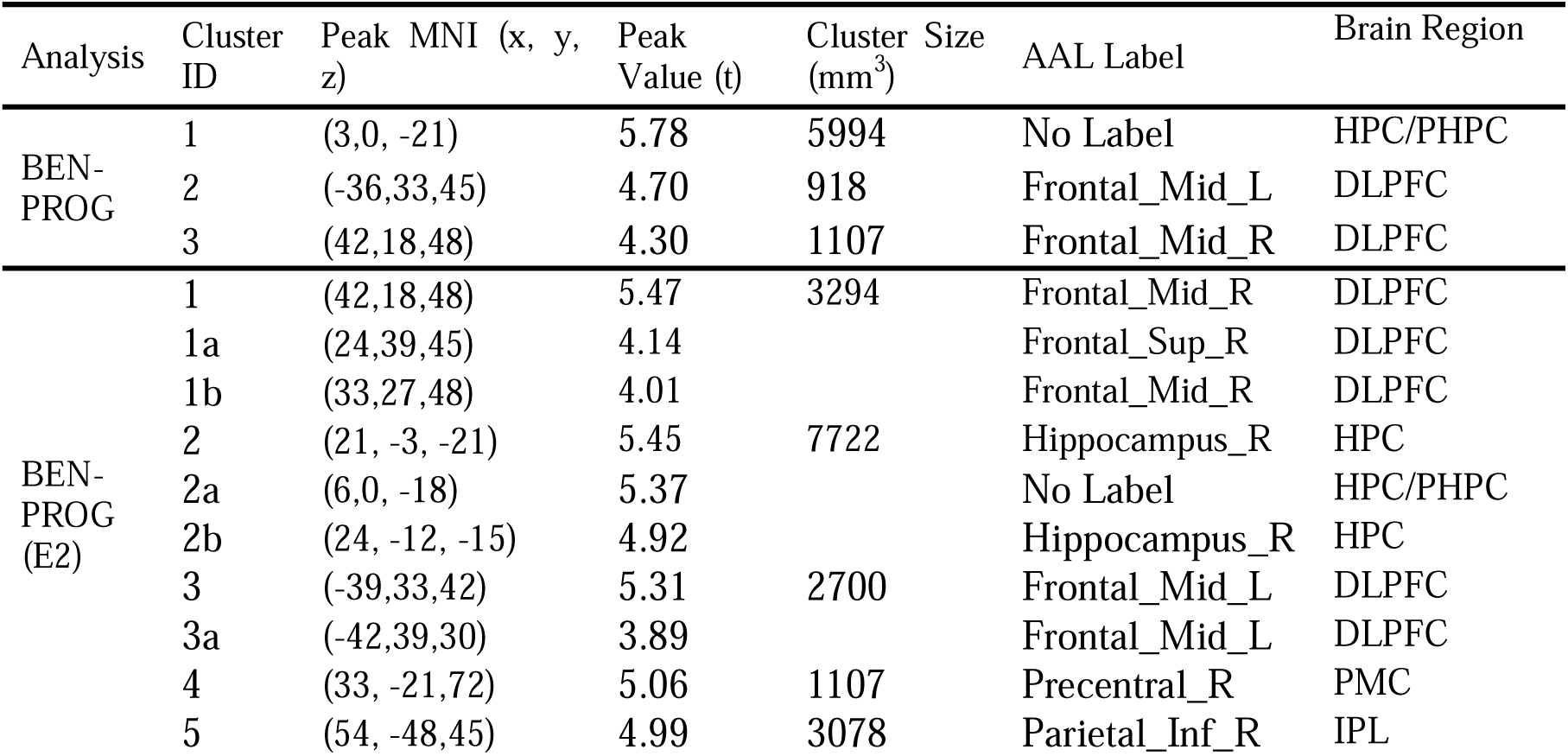
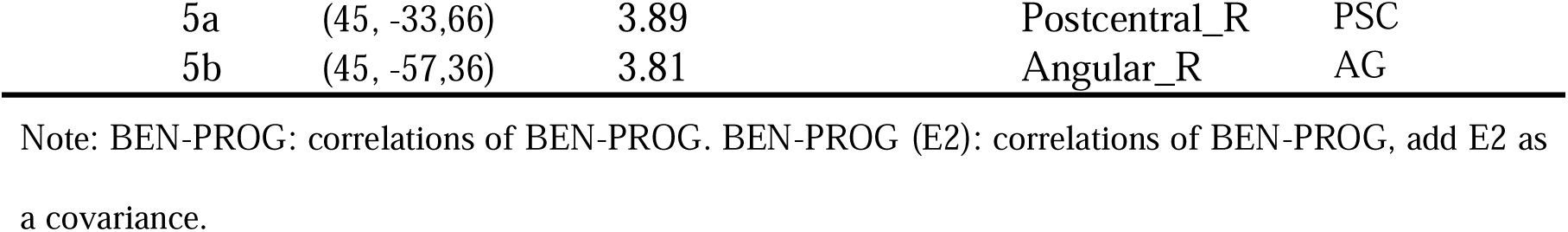
Clusters size table for correlation of BEN with ovarian hormones.

### 2.5 Correlation of BEN with BIS/BAS

The correlation between BEN and BAS-drive revealed a significant positive association in the left IFG and left DLPFC (Fig 2a). Upon adding PROG as a covariate, the previously positive correlation in the left DLPFC disappeared, but a negative correlation emerged in the medial temporal lobe (MTL) (Fig 2b). Upon adding E2 as a covariate, a negative correlation emerged in the medial temporal lobe (MTL) (Fig 2c). When both PROG and E2 were added as covariates, a similar pattern was observed when only PROG was included (Fig 2d). No significant correlations were detected between BEN and other BIS/BAS subscales. Table 3 provides information on clusters exceeding the threshold (voxel-wise p < 0.001, cluster size ≥ 810 mm^3^), including peak MNI coordinates, cluster size, label on the AAL atlas, and brain regions for correlation analysis of BEN with BIS/BAS.

**Fig 2.**
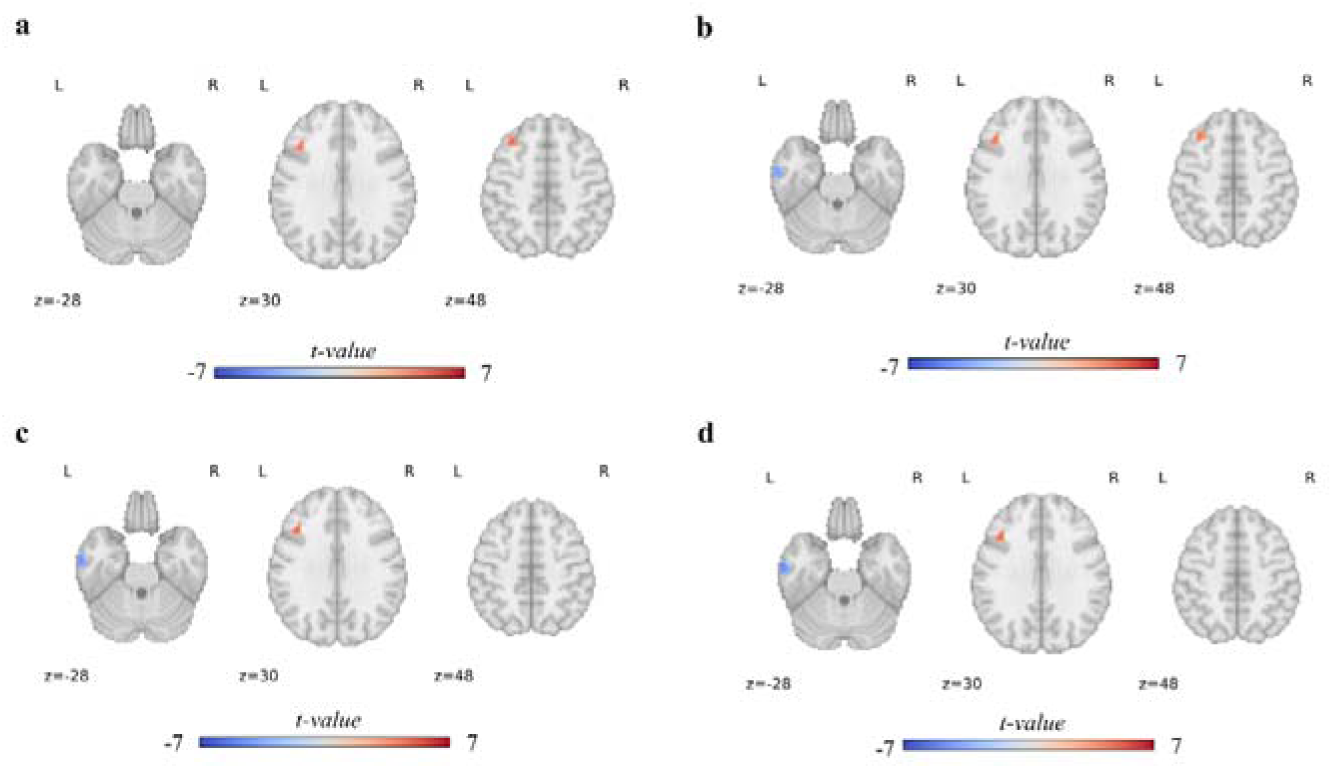
The correlations of BEN-BAS-drive, (a) without covariances, (b) add E2 as a covariance, (c) add PROG as a covariance, (d) add PROG and E2 as covariances. The color bar indicates the t value, the red indicates a positive correlation and the blue indicates a negative correlation between BEN and BAS-drive. The number beneath each slice indicates its location along the z-axis in the MNI space, and L means left hemisphere, and R means right hemisphere.

**Table 3.**
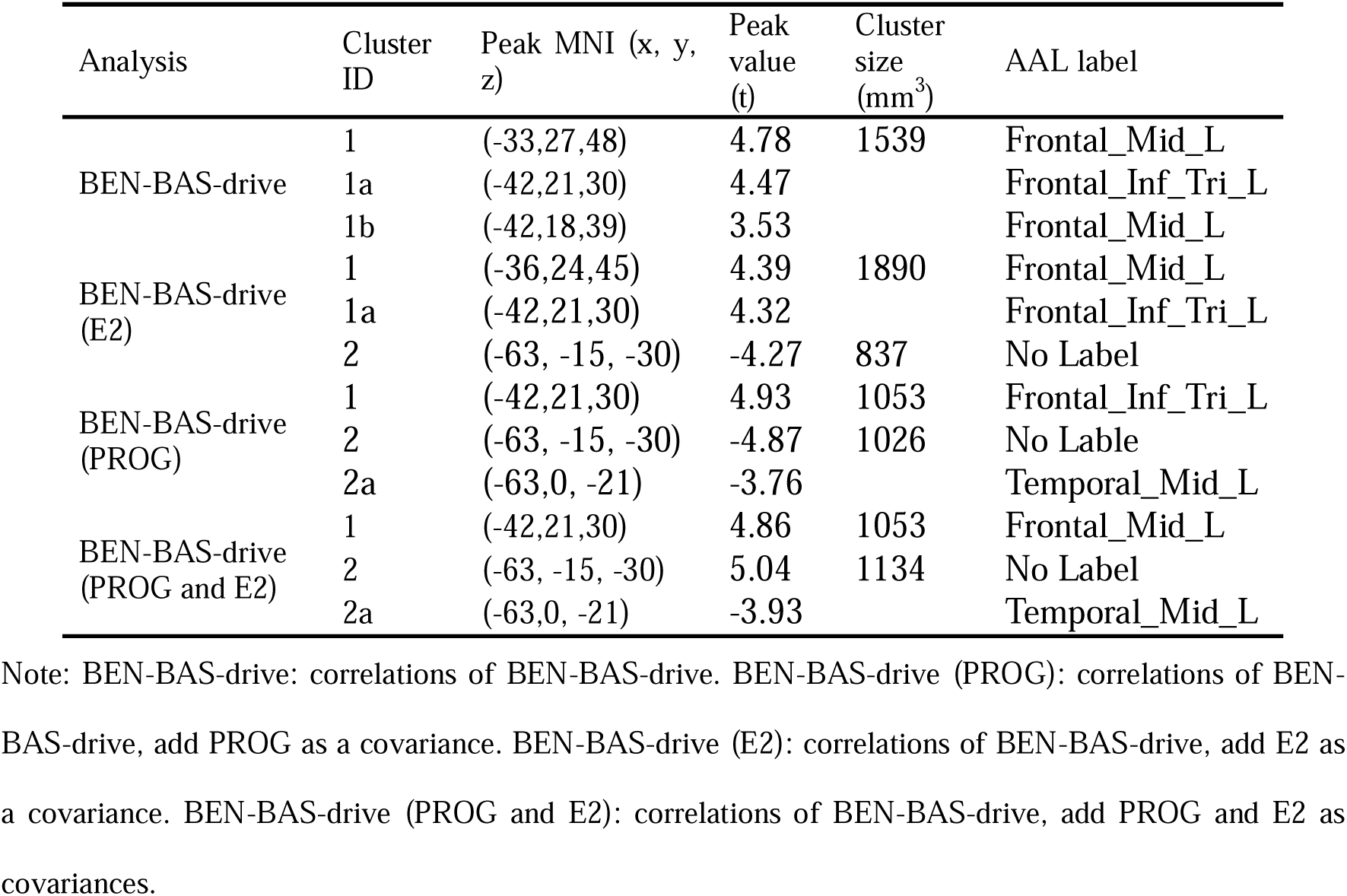
Clusters size table for correlation of BEN with BIS/BAS.

### 2.6 Correlation of DLPFC FC with BIS/BAS

The preceding results all demonstrate significant correlations between BEN and PROG, as well as between BAS-drive, within the left DLPFC. To further investigate the potential network connections underlying the associations between ovarian hormones and BAS, we also computed whole-brain functional connectivity using the DLPFC as a seed point. Please refer to the Methods section for specific details on the methodology.

The correlation between DLPFC FC and BAS-drive revealed a significant negative association in the left IPL, after adding E2 as a covariate, there were no noticeable changes in the results. However, when either adding PROG or including both E2 and PROG as covariates, no significant correlations were found between DLPFC FC and BAS-drive (Fig 3a). There exists a significant negative correlation between DLPFC FC and BAS-fun-seeking in the right lateral orbitofrontal cortex (LOFC), which is not influenced by ovarian hormones (Fig 3b). There is a significant negative correlation between DLPFC FC and BAS-rewards in the right AG, which is similarly unaffected by ovarian hormones (Fig 3c). Table 4 provides information on clusters exceeding the threshold (voxel-wise p < 0.001, cluster size ≥ 810 mm^3^), including peak MNI coordinates, cluster size, label on the AAL atlas, and brain regions for correlation analysis of DLPFC FC with BAS.

**Fig 3.**
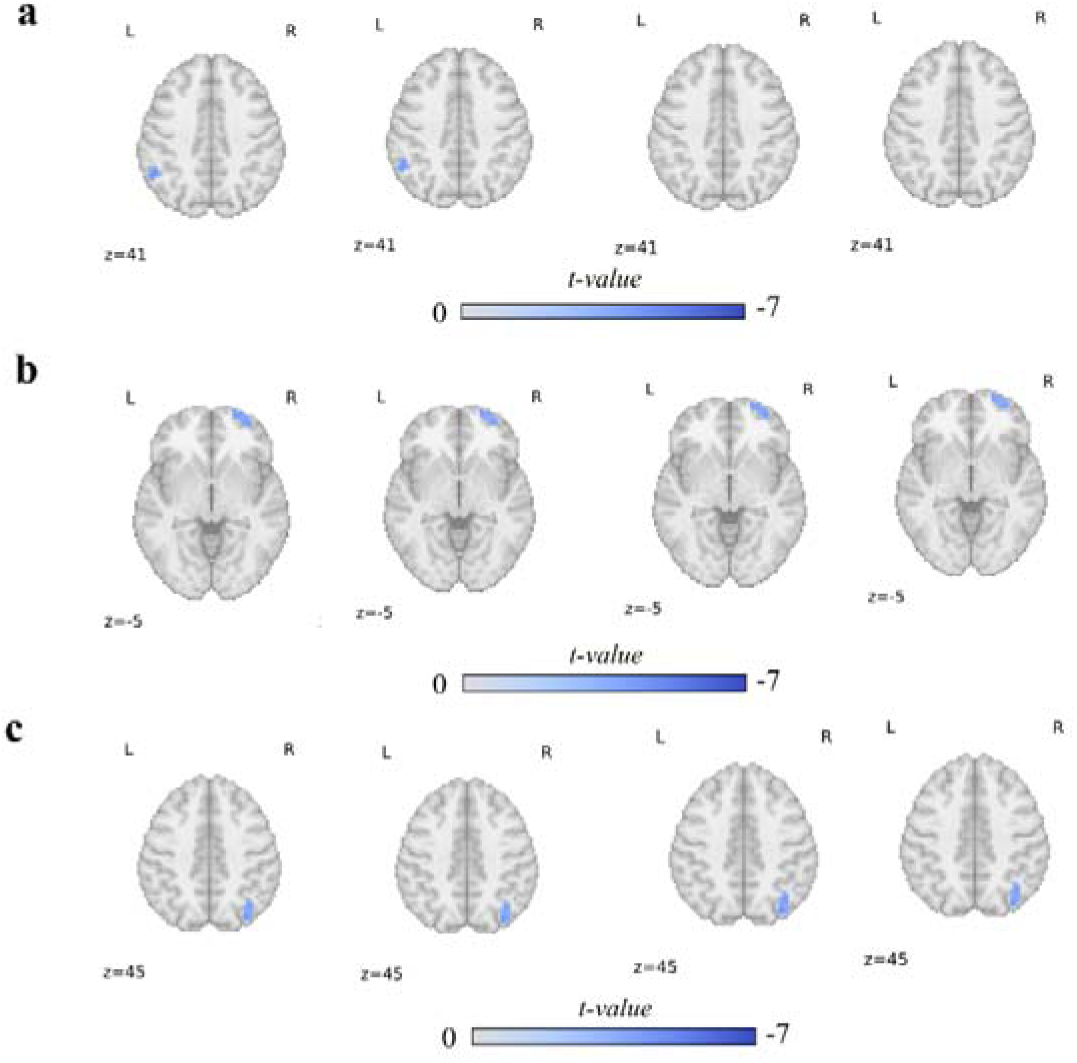
The correlations of DLPFC FC-BAS. (a) The correlations of DLPFC FC-BAS-drive, from left to right, the sequence is no covariates, adding E2 as a covariate, adding PROG as a covariate, and adding both E2 and PROG as covariates. (b) In the correlations of DLPFC FC-BAS-fun-seeking, from left to right, the sequence has no covariates, adding E2 as a covariate, adding PROG as a covariate, and adding both E2 and PROG as covariates. (c) The correlations of DLPFC FC-BAS-rewards, from left to right, the sequence is no covariates, adding E2 as a covariate, adding PROG as a covariate, and adding both E2 and PROG as covariates. The color bar indicates the t value, and the blue indicates a negative correlation between DLPFC FC and BAS. The number beneath each slice indicates its location along the z-axis in the MNI space, and L means left hemisphere, and R means right hemisphere.

**Table 4.** Clusters size table for correlation of BEN with BIS/BAS.

| Analysis | Cluster ID | Peak MNI (x, y, z) | Peak value (t) | Cluster size (mm<sup>3</sup>) | AAL label |
| --- | --- | --- | --- | --- | --- |
| FC-BAS-drive | 1 | (-54, -54,42) | 4.60 | 972 | Parietal_Inf_L |
| FC-BAS-drive (E2) | 1 | (-54, -54,42) | 4.49 | 837 | Parietal_Inf_L |
| FC-BAS-fun-seeking | 1 | (33,60,0) | 4.49 | 2295 | Frontal_Mid_R |
|  | 1a | (24,66, -6) | 4.47 |  | Frontal_Sup_Orb_R |
| FC-BAS-fun-seeking (E2) | 1 | (24,66, -6) | 4.31 | 1944 | Frontal_Sup_Orb_R |
|  | 1a | (33,60,0) | 4.26 |  | Frontal_Mid_R |
| FC-BAS-fun-seeking (PROG) | 1 | (24,66, -6) | 4.44 | 2403 | Frontal_Sup_Orb_R |
|  | 1a | (33,60,0) | 4.44 |  | Frontal_Mid_R |
| FC-BAS-fun-seeking (E2 and PROG) | 1 | (24,66, -6) | 4.24 | 1917 | Frontal_Sup_Orb_R |
|  | 1a | (33,60,0) | 4.20 |  | Frontal_Mid_R |
| FC-BAS-rewards | 1 | (33, -75,45) | 4.56 | 1917 | Occipital_Sup_R |
|  | 1a | (36, -63,39) | 4.20 |  | Angular_R |
|  | 1b | (51, -54,39) | 4.00 |  | Angular_R |
|  | 1c | (42, -57,39) | 3.93 |  | Angular_R |
| FC-BAS-rewards (E2) | 1 | (33, -75,45) | 4.51 | 1917 | Occipital_Sup_R |
|  | 1a | (36, -63,42) | 4.18 |  | Angular_R |
|  | 1b | (51, -54,39) | 4.10 |  | Angular_R |
| FC-BAS-rewards (PROG) | 1 | (33, -75,45) | 4.57 | 2025 | Occipital_Sup_R |
|  | 1a | (36, -63,45) | 4.23 |  | Angular_R |
|  | 1b | (42, -57,39) | 4.10 |  | Angular_R |
|  | 1c | (51, -54,39) | 3.91 |  | Angular_R |
| FC-BAS-rewards (E2 and PROG) | 1 | (33, -75,45) | 4.51 | 1998 | Occipital_Sup_R |
|  | 1a | (36, -63,42) | 4.26 |  | Angular_R |
|  | 1b | (42, -57,39) | 4.04 |  | Angular_R |
|  | 1c | (51, -54,39) | 3.92 |  | Angular_R |

Note: BEN-BAS-drive: correlations of BEN-BAS-drive. BEN-BAS-drive (PROG): correlations of BEN-BAS-drive, add PROG as a covariance. BEN-BAS-drive (E2): correlations of BEN-BAS-drive, add E2 as a covariance. BEN-BAS-drive (PROG and E2): correlations of BEN-BAS-drive, add PROG and E2 as covariances.

### 2.7 Correlation of hormones with BIS/BAS scoring, DLPFC BEN and DLPFC FC

The mean BEN values were extracted from the DLPFC ROI for each participant, while mean DLPFC FC z values were extracted from the region (IPL) where a significant correlation between DLPFC FC and BAS-drive was observed after regressing out E2 for each participant. Please refer to the Methods section for specific details on the methodology. Subsequently, correlations were conducted between age, mean FD, ovarian hormones, BIS/BAS scores, DLPFC BEN, and DLPFC FC.

The analysis results indicated no significant correlations between age and mean FD with any other variables. Significant positive correlations were found among the subscales of the BAS. Additionally, significant positive correlations were observed between PROG, E2, and DLPFC FC, while a significant negative correlation was noted between DLPFC BEN and BAS-d. Furthermore, a significant negative correlation was identified between DLPFC BEN and DLPFC FC (Fig 4).

**Fig 4.**
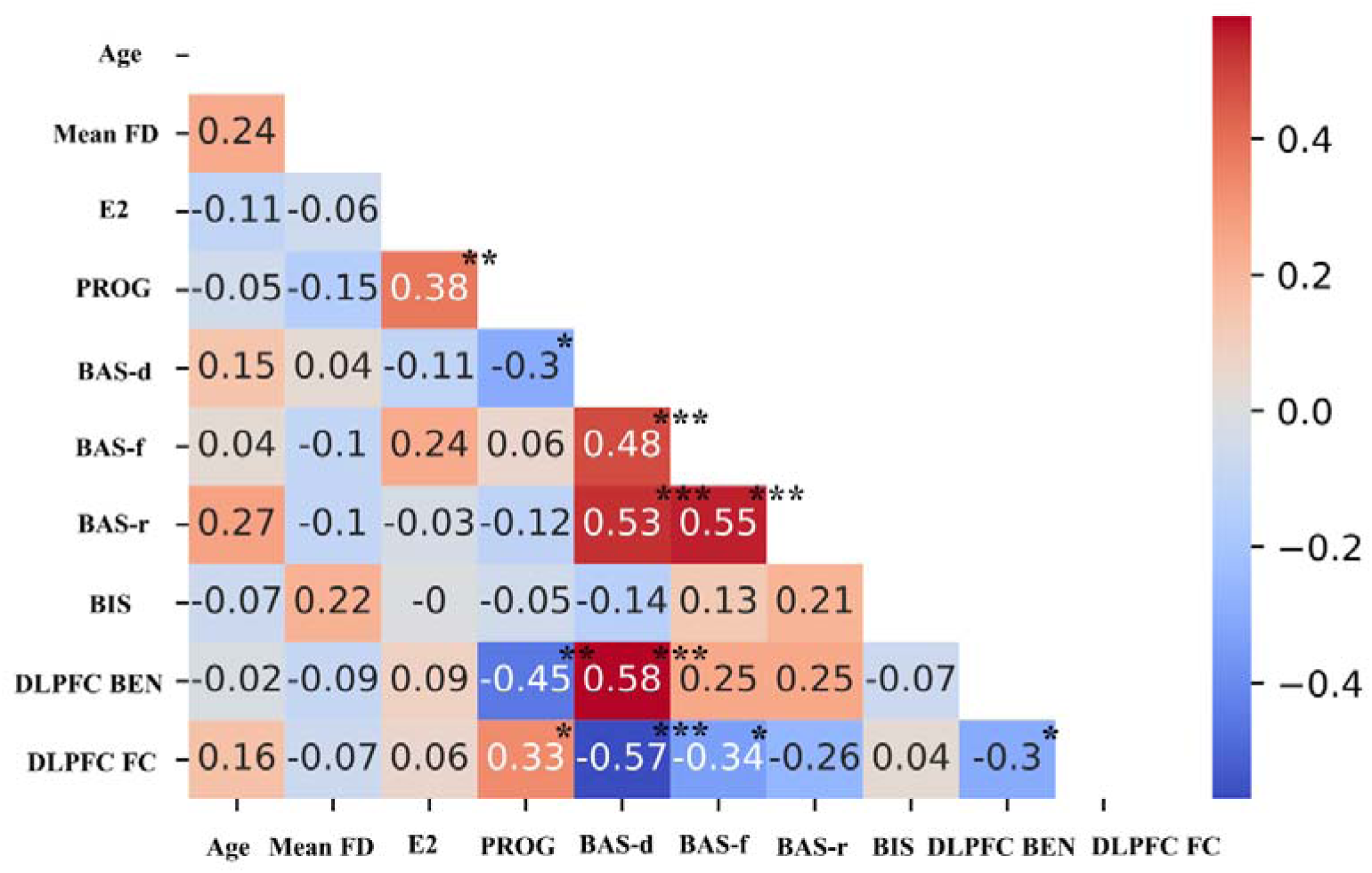
Heatmap of the correlation coefficient for ovarian hormones, BIS/BAS scoring, DLPFC BEN, and DLPFC FC. The colorbar indicates the r-value, the hot color indicates a positive correlation, and the blue color indicates a negative correlation. * p<0.05, **p<0.01, ***p<0.001.

### 2.8 Mediation Models Analysis

Based on the correlation analysis results, we observed significant pairwise correlations between PROG, BAS-d, DLPFC BEN, and DLPFC FC. To further explore their interrelationships, we constructed two mediation models: a chain mediation model and a parallel mediation model. Please refer to the Methods section for specific details on the methodology.

For the chain mediation model, we hypothesized that PROG affects DLPFC FC through its influence on DLPFC BEN, ultimately affecting BAS-d. However, our model did not hold (d21 = −0.006, p > 0.05, Fig 5a). For the parallel mediation model, we hypothesized that PROG affects BAS-d by separately influencing DLPFC BEN and DLPFC FC. However, the results indicated that only PROG’s influence on BAS-d through the mediator DLPFC BEN was significant, while DLPFC FC did not mediate the relationship between PROG and BAS- d (Fig 5b)

**Fig 5.**
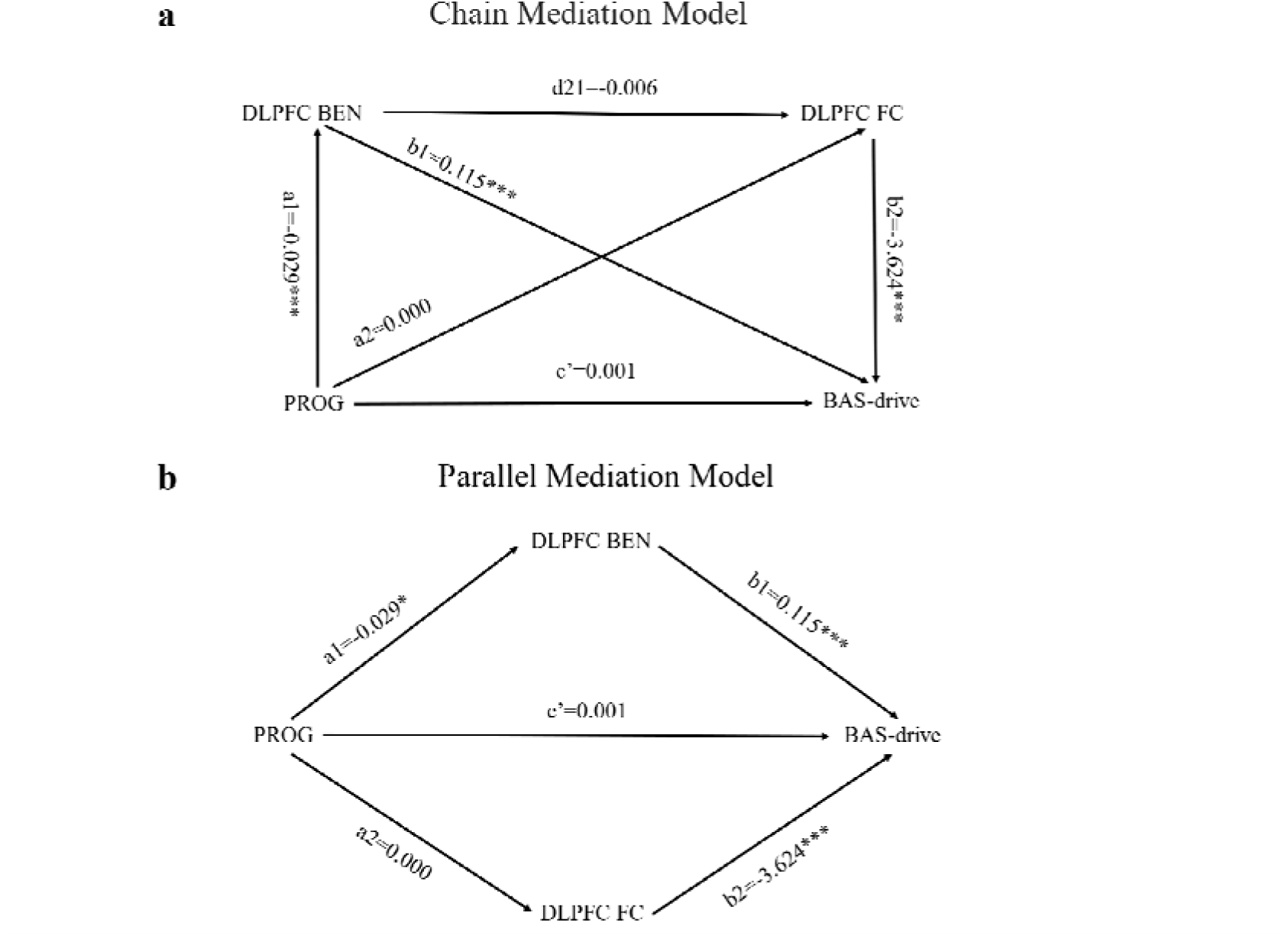
Mediation models analysis. (a) Chain mediation model; (b) Parallel mediation model. * p<0.05, **p<0.01, ***p<0.001.

## 3 Discussion

We first confirmed the relationship between BEN and ovarian hormones, the results showed that BEN and PROG had a significant negative correlation in DLPFC, IPL, AG, AMY, HPC, PHPC, and sensorimotor cortex (SMC) including PSC and PMC. The correlation analysis of BEN and BAS showed that BEN and BAS-drive found a significant positive correlation in the left DLPFC and IFG and a significant negative correlation in the MTL. DLPFC FC is significantly negatively correlated with BAS-drive in the IPL region, with BAS-fun-seeking in the right LOFC, and with BAS-rewards in the right AG. DLPFC FC is significantly positively correlated with PROG in the IPL. Furthermore, PROG influences BAS-drive through left DLPFC BEN.

### 3.1 The relationship of BEN with ovarian hormones and BIS/BAS

DLPFC and IPL are the core regions in the FPN, a crucial brain network for cognitive task switching, response inhibition, and executive control [57–59]. The negative correlation between PROG and BEN in this network suggests that higher levels of PROG are associated with greater neural activity coherence in FPN, potentially indicating better executive function. This is further supported by DLPFC FC, where there is a significant positive correlation between DLPFC FC and PROG in IPL. The studies by us and others together support this viewpoint, showing that lower BEN in the frontal and parietal cortex are associated with better fluid intelligence and functional task performance [18], while PROG has been shown to enhance executive control functions [36, 60–62].

AMY is a key brain region for emotion regulation [63, 64], while PHPC and HPC are known to be involved in memory and learning [65, 66], as part of the limbic system implicated in reward, motivation, addiction, memory, and emotion [67–69]. The studies consistently found that during the LP, when PROG levels are high, there is an increased AMY response to negative emotional stimuli [70, 71]. There are conflicting results regarding the role of PROG in memory, with the study by Ertman et al. showing positive correlations with recall and recognition of memory for emotional stimuli [72], while studies by Van Wingen et al. indicate that PROG impairs memory [73, 74]. This may be attributed to the bottom-up regulation between the limbic system and DLPFC by PROG. A recent study suggested that negative emotion processing activates the limbic system and PROG modulates more bottom-up brain activation during both positive and negative emotion processing [37]. This finding aligns with our analysis of DLPFC FC, wherein DLPFC-IPL FC positively correlates with PROG, suggesting that stronger executive control functions are associated with PROG. However, no significant negative correlation was observed between DLPFC FC and PROG in the limbic system, even when the threshold was relaxed, a positive correlation was observed in the left AMY.

The SMC plays a major role in controlling how the body responds to external stimuli [75]. A study using eigenvector centrality mapping revealed a significant positive correlation between PROG and eigenvector centrality in the bilateral DLPFC and bilateral SMC on the whole brain level, along with a PROG-modulated increase in FC of both bilateral DLPFC and bilateral SMC with the HP [76].

### 3.2 The relationship of BEN, DLPFC FC, PROG, and BIS/BAS

There is a significant positive correlation between BEN and BAS-drive in the left DLPFC and IFG, while a negative correlation is observed in the MTL. DLPFC FC analysis showed a negative correlation between DLPFC FC and BAS-drive in the left IPL, a significant negative correlation between DLPFC FC and BAS-fun-seeking in the right LOFC, and a significant negative correlation between DLPFC FC and BAS-rewards in the right AG. On the other hand, PROG and BAS-drive demonstrated a positive correlation. Through mediation analysis, it is confirmed that PROG influences BAS drive through the mediator DLPFC BEN.

BAS-drive is associated with reward and impulsivity addiction, with a higher BAS-drive leading to increased impulsive behaviors and substance abuse. Both DLPFC and IFG are related to executive control. The positive correlation implies that higher BEN in DLPFC and IFG is associated with an increased risk of impulsivity and substance abuse. This aligns with our previous research indicating a higher risk of substance abuse in adolescents with higher BEN in the FPN [56], as well as our findings showing a higher BEN in the DLPFC among individuals with substance use disorders [22], and consistent results of higher BEN in the DLPFC among smokers [25]. However, when PROG was added as a covariate, this positive correlation between DLPFC BEN and BAS-drive disappeared, while a significant negative correlation between PROG and BAS-drive was observed. Through mediation analysis, it has been confirmed for the first time that PROG influences the behavioral activation system (BAS)-drive via dorsolateral prefrontal cortex (DLPFC) executive network (BEN). In essence, PROG enhances executive function in the DLPFC, thereby reducing BAS-drive. Other studies consistently demonstrated that PROG could attenuate addictive behaviors [77–79]. The MTL is responsible for facial perception, emotional perception, and semantic understanding [80–82]. The negative correlation between BEN and BAS-drive in the MTL suggests that individuals with higher BAS-drive, indicative of the greater desire for rewards, positive affect, and motivational drive, demonstrate faster recognition of language information and perception of emotional cues.

The results of DLPFC FC do not support the hypothesis that PROG influences distant brain regions through its effects on local brain activity (DLPFC BEN) to impact BAS-drive. Instead, the negative correlation between DLPFC and left IPL FC independently correlates with BAS-drive, indicating that the enhanced FC between DLPFC and left IPL is associated with higher executive function and inhibitory control, which aligns with the impulsivity represented by BAS-drive. Furthermore, DLPFC FC and BAS-fun-seeking in the right LOFC exhibit a significant negative correlation. Since LOFC is associated with reward processing, the enhanced DLPFC-LOFC FC suggests an improved control function over rewards, consistent with the sensation-seeking represented by BAS-fun-seeking. Additionally, the negative correlation between DLPFC FC and BAS-rewards in the right AG suggests a consistent relationship between higher executive function and inhibitory control desires with the reward-seeking represented by BAS-rewards.

## 4 Conclusion

In conclusion, BEN demonstrates sensitivity to ovarian hormones, serving as a monitor for the neural effects of these hormones, and the relationship between BEN and BAS-drive is also influenced by ovarian hormones that left DLPFC mediates the relationship between PROG and BAS-drive. Our research also provides neurobiological support for emphasizing the role of women’s endogenous hormonal concentrations in emotion and cognition.

## 5 Methods

### 5.1 Dataset

The data of the study from OpenNeuro ds003114 that released by Jia-Xi Wang et al (https://openneuro.org/datasets/ds003114/versions/1.0.0). The dataset includes 49 healthy participants, ages from 19 to 28 years old (mean age = 22.77 years, standard deviation (SD) = 2.35). The backward counting method was used to predict each participant’s late FP, mid-LP, and next menstrual onset. The late FP included the period of 14 to 16 days before a woman’s next predicted menstrual onset, and the mid-LP included the period of 6 to 8 days before her next predicted menstrual onset. 25 women in the late FP with a mean age of 22.54 years old (SD = 2.18) and 24 women in their mid-LP with a mean age of 23.04 years (SD = 2.54) were subjected to the resting-state fMRI. The testing order was randomized across participants and phases.

A saliva sample was obtained from each participant immediately before scanning to quire E2 and PROG levels. Participants completed a Chinese version of the BIS/BAS Scale before the resting-state scanning. The BIS subscale relates to the anticipation of punishment and avoidance of cues to negative outcomes. The BAS is divided into three subscales: a positive response to the occurrence or anticipation of reward (BAS-reward); a persistent pursuit of desired goals (BAS-drive); and a desire for new rewards and willingness to approach a potentially rewarding event on the spur of the moment (BAS-fun seeking).

The high-resolution structural images and rs-fMRI images were acquired on a 3-T Siemens scanner. The rs-fMRI images were acquired with a gradient echo-planar imaging sequence with repetition time (TR) =2000 ms, echo time (TE) = 30 ms, field of view (FOV) =384 mm, 3 mm × 3 mm × 3.5 mm voxel size and 33 slices and participants eyes open during scanning which lasted about 8 minute and total 240 volumes were acquired. A high-resolution structural image was acquired with a T1-weighted, multiplanar reconstruction sequence (TR = 2530 ms, TE = 2.98 ms, FOV = 256 mm, 1 mm × 1 mm × 1 mm voxel size, and 192 slices).

For more detailed participants’ information, hormone assays, BIS/BAS scale, and MRI acquisition parameters can be found in the original article from the dataset [46].

### 5.2 MRI preprocessing

The MRIs were preprocessing using fmriprep [83] (version=23.1.4) (https://pypi.org/project/fmriprep-docker/), which is based on Nipype (version=1.8.6) [84], is containerized to doc ker (version=24.0.7) (https://www.docker.com/) under Ubuntu (version=22.04.3 LTS) (https://releases.ubuntu.com/jammy/), then denoising using based-Nilearn (version=0.10.2) (https://nilearn.github.io/stable/index.html) customed python (version=3.10) (https://www.python.org/) scripts.

The structural images were skull-stripped and segmented into gray matter (GM), white matter (WM), and cerebrospinal fluid (CSF) using FAST [85], then normalized to MNI space. The functional images underwent preprocessing through the following steps: (1) Slice-time corrected to 0.96s (0.5 of slice acquisition range 0s −1.92s) using 3dTshift from AFNI [86]. (2) Head motion correction was performed using mcflirt [87]. (3) The functional reference was t hen co-registered to the T1w reference using mri_coreg followed by flirt [88] and framewise displacement (FD) [89], and DVARS were calculated. The average signals were extracted wit hin the GM, CSF, and WM. (4) The functional images were resampled into MNI space using antsApplyTransforms (ANTs). (5) Detrending and temporal bandpass filtering (0.009–0.08 H z) was performed. (6) Nuisance components were regressed out including the six motion para meters, FD, std_dvars, rmsd, tcompcor, and first three componence of c_comp_cor, as well as the average WM signals and average CSF signals. (7) Smoothing with an isotropic Gaussian kernel (FWHM□=□6 mm). For more detailed preprocessing details, please refer to [31].

### 5.3 BEN mapping calculation

BEN was calculated at each voxel of the preprocessed functional images by the BEN mapping toolbox (BENtbx) [10]. The toolbox can be found at https://www.cfn.upenn.edu/zewang/BENtbx.php and https://github.com/zewangnew/BENtbx. In this study, the window length was set to 3 and the cut-off threshold was set to 0.6. The first four volumes were discarded for signal stability and to reduce variability due to noise, BEN maps were smoothed with an isotropic Gaussian kernel with FWHM□=□8 mm. More details of BEN calculation can be found in the original BEN paper (Wang et al., 2014) or in our previous studies [16, 18, 20, 21, 90].

### 5.4 DLPFC FC calculation

We selected the overlapping regions where BEN and PROG as well as BAS-drive showed significant association as a region of interest (ROI). The ROI was binarized to create a mask and utilized as a seed for whole-brain functional connectivity (FC) analysis. The purpose of this analysis is to determine whether the DLPFC BEN influences distant effects of DLPFC and subsequently, whether the DLPFC FC is associated with ovarian hormones and BIS/BAS.

The specific seed-based FC calculation was as follows: FC maps were calculated by customed Python scripts based on Nilearn (https://nilearn.github.io/stable/index.html) using after preprocessed rs-fMRI images. The mean value of the rs-fMRI time series was extracted from the seed and correlated to the time series of all voxels in the whole brain. Pearson’s correlation coefficient was measured as the amplitude of FC and Fisher’s z-transformed to improve the normality before performing group-level analysis.

### 5.5 Statistical analysis

All statistical analyses were performed using customized Python scripts. The statistical significance level for the analysis of demographic information, hormone assays, and BIS/BAS was set at *P* < 0.05. Significance thresholds were set at voxel-wise *P* < 0.001 with a cluster size > 540 mm^3^ (30 voxels) for BEN differences between LP and FP, as well as the analysis of the correlation of BEN with hormones and BIS/BAS.

#### 5.5.1 Demographics and Hormone assays and BIS/BAS scoring

Use NumPy (https://numpy.org/) to calculate the mean and SD of the ages of all subjects, as well as for the corresponding FP and LP. Apply a two-sample t-test to determine if there are significant differences in age and mean FD between LP and FP. E2, PROG, BIS, BAS-drive, BAS-reward, and BAS-fun seeking differences between FP and LP were tested using a two-sample t-test.

#### 5.5.2 BEN differences between LP and FP

Use a two-sample t-test to determine BEN differences during LP and FP, with age, and mean FD as confounds. To determine the impact of PROG and E2, we then also included PROG and E2 separately or together as covariances into the general linear model (GLM) to see the impact on FP and LP.

#### 5.5.3 Analysis of the correlation of BEN with Hormones and BIS/BAS

GLM was used to evaluate the correlation between BEN with hormones, age, and mean FD as confounds. To determine the impact of PROG and E2, we then also included PROG or E2 separately or together as covariances into the GLM to see the impact on E2 or PROG. GLM was used to evaluate the correlation between BEN with BIS/BAS, with age, and mean FD as confounds. To determine the impact of PROG and E2, we then also included PROG and E2 separately or together as covariances into the GLM to see the impact on BIS/BAS.

#### 5.5.4 Analysis of correlation of DLPFC FC with Hormones and BIS/BAS

DLPFC performs analysis like BEN with Hormones and BIS/BAS in 5.5.3.

#### 5.5.5 Analysis of correlation of DLPFC BEN with DLPFC FC, Hormone assays, and BIS/BAS scoring

Mean BEN values were extracted from the DLPFC ROI for each participant, while mean DLPFC FC z values were extracted from the region (IPL) where a significant correlation between DLPFC FC and BAS-drive was observed after regressing out E2 (Fig 3a).

#### 5.5.6 Mediation Models Analysis

According to the results of the correlation analysis (Fig 4), significant correlations were found among PROG, BAS-drive, DLPFC BEN, and DLPFC FC. Based on these results, we constructed two mediation models. The first model (Fig 5a) is a chain mediation model, in which we hypothesized that PROG (independent variable) affects DLPFC BEN (mediator variable 1), which in turn influences DLPFC FC (mediator variable 2), ultimately affecting BAS-drive (dependent variable). The second model (Fig 5b) is a parallel mediation model, in which we hypothesized that PROG (independent variable) influences BAS-drive (dependent variable) through DLPFC BEN (mediator variable 1) or DLPFC FC (mediator variable 2). Statistical analysis was performed with the “lavaan” package (https://lavaan.ugent.be/tutorial/mediation.html) [91] in R version 4.1.2 (https://www.r-project.org/). The R code including model setup is available at https://github.com/donghui1119/Brain_Entropy_Project/tree/main/Hormones (up on publication of the manuscript).

## Acknowledgments

We thank Jia-Xi Wang et al for releasing their dataset.

## Data and code availability

All raw data are available at OpenNeuro ds003114 (https://openneuro.org/datasets/ds003114/versions/1.0.0).

BENtbx is available at https://www.cfn.upenn.edu/zewang/BENtbx.php.

Lavaan is available at https://lavaan.ugent.be/

Customed codes and further updates related to the study will be available at https://github.com/donghui1119/Brain_Entropy_Project/tree/main/Hormones (upon publication of the m anuscript).

## CRediT authorship contribution statement

Dong-Hui Song: conceptualization, data curation, data analysis, visualization, manuscript drafting, and editing. Ze Wang: conceptualization, manuscript editing, supervision, project a dministration.

## Notes

https://openneuro.org/datasets/ds003114/versions/1.0.0

### Competing Interest Statement

The authors have declared no competing interest.

